# Negative-Valence Neurons in the Larval Zebrafish Pallium

**DOI:** 10.1101/2025.11.21.689820

**Authors:** Colton D. Smith, Zhuowei Du, William P. Dempsey, Scott E. Fraser, Thai V. Truong, Don B. Arnold

## Abstract

An organism’s survival depends on the rapid classification of sensory events as either harmful or beneficial. In mammals, this computation is partially performed by neurons in the amygdala that respond to signals with negative or positive valence. The larval zebrafish pallium is thought to contain homologs of the mammalian amygdala, isocortex, and hippocampus; however, the signals encoded by pallial neurons remain largely uncharacterized. Using two-photon light-sheet microscopy to image 7–9-day-old zebrafish that express pan-neuronal GCaMP6s, we recorded calcium dynamics throughout the brain while presenting a panel of strongly aversive stimuli including infrared heat, electric shock, and a whole-field looming shadow, together with milder threats including vibration, loud sound, light transitions, and a partial-looming stimulus. A compact cluster of neurons in the rostrolateral dorsal pallium (Rl) responded vigorously to strongly noxious and fully looming stimuli, but not to the milder cues. In contrast, neurons in the ventromedial pallium and habenula responded to all stimuli tested. Rl neurons are characterized by high *tiam2a* expression, suggesting they can be genetically accessed. Thus, our results identify a locus of negative-valence neurons in the teleost pallium.

## Introduction

To evaluate potential threats, organisms must predict future events based on sensory cues (Herberholz and Marquart, 2012). For example, when a predator approaches a fish, sudden changes in water pressure activate its lateral line system, which, combined with visual cues, allows it to predict that danger is imminent and swiftly evade the threat (Olszewski et al., 2012). How fish decide to flee, or, conversely, to mate or feed, remains poorly understood. In mammals, several lines of evidence suggest that such decisions involve the basolateral amygdala, where distinct ensembles encode negative and positive value (Beyeler et al., 2018; Grewe et al., 2017; Namburi et al., 2015; Tye, 2018; Xiu et al., 2014). Lesions or chemogenetic silencing of the amygdala blunt defensive behaviors and block fear conditioning (Bechara et al., 2003; Corder et al., 2019; Downer, 1961; Feinstein et al., 2011; LeDoux et al., 1988; Phillips and LeDoux, 1992). The amygdala is also a locus of synaptic change following fear learning (Clem and Huganir, 2010; Goosens and Maren, 2002; Nabavi et al., 2014; Phillips and LeDoux, 1992; Quirk et al., 1995; Rogan et al., 1997). Finally, neurons in the basolateral amygdala project differentially depending on their valence, with negative-valence neurons projecting to the medial central amygdala and neighboring positive-valence neurons projecting to the nucleus accumbens, consistent with activation of the two sets of neurons causing distinct downstream effects (Beyeler et al., 2018; Fadok et al., 2017; Penzo et al., 2014; Tye et al., 2011).

The larval zebrafish (*Danio rerio*) is an excellent model system for studying sensory processing because its transparent brain is amenable to whole-brain imaging paradigms that require low scattering, such as selective plane illumination microscopy (SPIM; Ahrens et al., 2013). The high speed of SPIM and its low toxicity enable 4-D imaging across large areas of the larval zebrafish brain (Vladimirov et al., 2018). Furthermore, a modified SPIM that uses two-photon (2P) excitation (Truong et al., 2011) significantly reduces toxicity and increases penetration relative to one-photon SPIM. Although its brain is smaller and less complex than those of many other vertebrates, the zebrafish can, even in its larval stage, learn to modify responses to environmental stimuli (Gerlai, 2011). In particular, zebrafish are amenable to classical conditioning, which in some cases depends on distinct regions in the telencephalon (Lal et al., 2018; M Portavella et al., 2004; Portavella et al., 2002; Ruhl et al., 2015). Previously, we showed that a well-defined area in the rostrolateral pallium (Rl) contains neurons that respond to aversive heating (Dempsey et al., 2022), suggesting that Rl, like the mammalian amygdala, may contain negative-valence neurons.

To characterize how neurons in the forebrain of a larval zebrafish respond to sensory stimuli of different valence, we mapped GCaMP (a genetically encoded Ca^2+^ indicator) fluorescence following exposure of the fish to stimuli ranging from highly threatening to mildly aversive. We found a tightly clustered group of neurons in Rl that responded vigorously to noxious or looming stimuli, but not to less intense negative-valence stimuli. Two additional areas, the ventromedial pallium (Vm) and the habenula (Hb), contained neurons that responded to both types of stimuli. Negative-valence neurons in Rl specifically expressed *tiam2a*, a homolog of a gene expressed in the mouse hippocampus. Thus, we have identified a set of negative-valence neurons within the zebrafish pallium that express a unique molecular marker.

## Results

### Noxious stimuli activate pallial neurons

To investigate how neurons in the forebrain respond to negative-valence stimuli, we created a transgenic zebrafish line, Tg[CAG_NRSE_:GCaMP6s], expressing GCaMP6s (Chen et al., 2013) driven by the chicken β-Actin CAG promoter (Quitschke et al., 1989) with an NRSE repressor element (Bergeron et al., 2012) added to diminish expression in non-neuronal cells. Wholemount immunostaining for GCaMP6s and the neuronal marker HuC (elav3) in the forebrain of a 7 dpf Tg[CAG_NRSE_:GCaMP6s] zebrafish shows almost complete overlap of the two labels, suggesting that virtually all neurons in the zebrafish pallium express GCaMP6s (***Figure 1-figure supplement 1***). Note that some skin cells expressed GCaMP6s and were excluded from all analyses.

**Figure 1.**
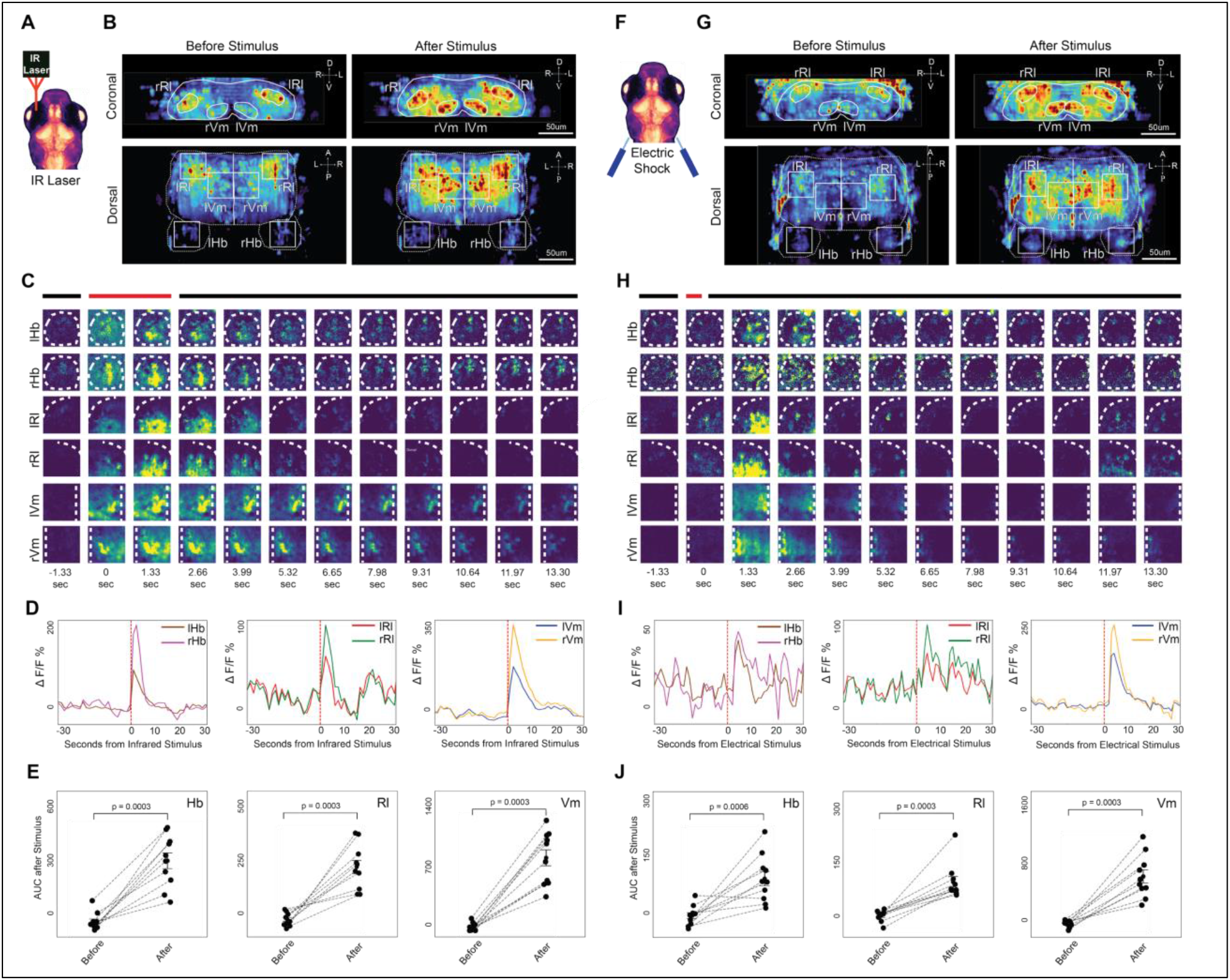
Neuronal activation in response to IR heating and electric shock. (**A**) Experimental setup with an infrared (IR) laser aimed at a 7 dpf larval zebrafish to deliver a heat stimulus. (**B**) Heat maps of GCaMP fluorescence from a representative fish averaged over 5 presentations of IR heating. Enhanced activity is displayed in rostrolateral (lRL, rRL) and ventromedial (lVm, rVm) areas of the pallium, and habenula (lHb, rHb) in coronal and dorsal views post-laser exposure. (**C**) Timelapse of GCaMP fluorescence heatmaps in response to exposure to the IR laser in the forebrain of a representative zebrafish. Each square corresponds to an area outlined in the dorsal view in (B). Brain regions where activity measurements were taken are outlined with dotted lines. Background activity was subtracted from all images. (**D**) Region-specific ΔF/F activity traces from a representative fish, averaged across 5 trials, display pronounced increases in all measured areas following IR laser exposure. (**E**) Analysis of the area under the curve (AUC; 5 frames, 6.65 seconds) ΔF/F for each fish for each brain region outlined in (**B**, **C**). Mean ± SEM for n=5 fish. Note that graphs contain measurements for the same region on both sides of the brain. (**F**) Illustration of electric shock delivery to a 7 dpf larval zebrafish. (**G**) Heat maps of GCaMP fluorescence from a representative fish averaged over 5 presentations of electric shock. Enhanced activity is displayed in rostrolateral (lRL, rRL) and ventromedial (lVm, rVm) areas of the pallium, and habenula (lHb, rHb) in coronal and dorsal views post-laser exposure. (**H**) Timelapse of GCaMP fluorescence heatmaps in response to exposure to electric shock in the forebrain of a representative zebrafish. Each square corresponds to an area outlined in the dorsal view in (G). Brain regions where activity measurements were taken are outlined with dotted lines. Background activity was subtracted from all images. (**I**) Region-specific ΔF/F activity traces from a representative fish, averaged across 5 trials, display pronounced increases in all measured areas following electric shock. (**J**) Analysis of the area under the curve (AUC; 5 frames, 6.65 seconds) ΔF/F for each fish for each brain region outlined in (G, H). Mean ± SEM for n=5 fish. One-tailed t-test was used for all statistical comparisons. Red dashed lines denote the onset of the stimulus. Abbreviations: A, anterior; AUC, area under the curve; D, dorsal; L, left; lHb, left habenula; lRl, left rostrolateral; lVm, left ventromedial; P, posterior; R, right; rHb, right habenula; rRl, right rostrolateral; rVm, right ventromedial; V, ventral. **Figure 1-source data 1.** Numerical source data for graphs in ***Figure 1D***, ***1E***, ***1I***, and ***1J***. **Figure1-figure supplement 1**. Tg[CAGNRSE::PGCaMP6s] expresses GCaMP6s in all neurons. **(A)** 7 dpf Tg[CAG_NRSE_::GCaMP6s] zebrafish dorsal pallium imaged for GCaMP6s. (**B**) Same zebrafish dorsal pallium immunostained for HuC, which labels all neurons. (**C**) Merge of (A) and (B) shows that essentially all cells labeled for HuC are also labeled for GCaMP6s. Arrowheads point to skin cells expressing GCaMP6s, which were excluded from all analyses. Scale bar 25 μm **Figure 1-figure supplement 2**. Two-photon light sheet microscopy setup for whole-brain functional imaging See (Keomanee-Dizon et al., 2020) for details on design, construction, and operation of the setup. Briefly, the mounted zebrafish sample (depicted as the orange ellipsoid at the center) was positioned inside an aqueous chamber, where excitation light sheets (depicted in red) illuminated the zebrafish’s brain from both the front and (left-lateral) side of the animal. Inset at lower right shows a picture of the mounted zebrafish, immobilized in 1.5% low-melt agarose, with its tail freed. Wide-field video recording from the bottom of the sample chamber provided readout of the zebrafish’s behavior through its tail movement. The excitation light sheets came from a femtosecond-pulsed near-infrared laser, with wavelength ∼920 nm, to generate two-photon-absorption fluorescence from the brain of the zebrafish. The fluorescence signal (depicted in green) was collected by the upright-oriented detection objective and imaged onto a recording camera (not shown). Physical masks made from black-anodized aluminum were positioned along the optical path of the light sheets to prevent direct illumination of the zebrafish eyes. Light from a continuous-wave near-infrared laser, wavelength ∼980 nm, was directed toward the zebrafish sample through the detection objective to provide an option for heating stimulation. **Figure 1-figure supplement 3. Recording areas** Recordings were conducted in 50 μm square areas. (**A**) The habenula recordings come from a superficial slice at 10 μm below the surface of the fish brain, 40 μm from the rostral edge, and adjacent to the most lateral edge of the brain. (**B**) The rostrolateral recordings were done at 30 μm below the surface of the brain, adjacent to its rostral edge, and roughly 10 μm from the midline. (**C**) The ventromedial recordings were done at 50 μm below the surface 40 μm from the rostral edge of the brain and adjacent to the midline.

To explore the pallial responses to negative-valence stimuli in real time, we imaged the Tg[CAG_NRSE_:GCaMP6s] zebrafish using a custom-built 2P SPIM (Keomanee-Dizon et al., 2020; ***Figure 1-figure supplement 2***). The 2P-SPIM microscope generates a 2P-excitation light sheet using ultrafast near-infrared laser pulses. This microscope combines non-linear excitation for high penetration depth with orthogonal light-sheet illumination for high acquisition speed and low photodamage (Truong et al., 2011). We immobilized 7–9 dpf fish by embedding them in agarose and placing them in a holder immediately adjacent to the SPIM’s two objectives (***Figure 1-figure supplement 2***).

In our initial studies, we exposed the fish to noxious infrared (IR) heating from a laser positioned over the zebrafish’s right eye (***Figure 1A, Figure 1-figure supplement 2***), which we had previously found to elicit a strong escape response consisting of a tail flick (Dempsey et al., 2022). The IR stimulus consisted of a series of five 2-second pulses with 2-minute intervals between. Before, during, and after the stimulus, the SPIM imaged a series of optical sections at 11 depths spanning 60 µm, acquired sequentially every 0.12 seconds, completing a stack every 1.33 seconds. We found that three discrete areas in each hemisphere of the zebrafish forebrain showed an overall increase in fluorescence (ΔF/F, see methods) in response to the IR stimulus (***Figure 1B***). These areas correspond to the Hb, Rl, and Vm areas (***Figure 1-figure supplement 3***). Time-lapse images of fluorescent signals (with background measured before the stimulus subtracted) show an abrupt increase shortly after the stimulus, with average ΔF/F increasing significantly in each of the six areas (***Figure 1C-E***, n = 5 fish). Note that because measurements of AUC for the same areas (Hb, Rl, or Vm) on the left vs. right were found not to be significantly different (p > 0.7, ANOVA, multiple comparisons), left and right regions were combined on the same graphs (***Figure 1E***).

To characterize the response of pallial and habenular neurons more fully, we exposed the fish to electric shock, another noxious stimulus. We used wires placed approximately 1 mm from the otic vesicles (the ears) to deliver 500 mA of current for 400 ms (***Figure 1F***). All brain regions that increased activity with IR stimulation also showed a qualitative increase in activity following the electric shock (***Figure 1G, H***). In addition, the average ΔF/F, measured after electric shock, was significantly higher than measurements before the shock (n = 5 fish, ***Figure 1I, J***). These results show that the Hb, Rl, and Vm areas of the pallium bilaterally respond to two different noxious stimuli.

To better understand activity at the level of individual neurons, we used the Ca^2+^ signal detection program CaImAn (Giovannucci et al., 2019) to extract neuronal responses from the raw fluorescence data (see methods). Because the CaImAn program cannot reliably distinguish signals from axons, dendrites, and somata, the signals may not represent distinct cells; thus, we refer to the traces produced by CaImAn as “neuronal components”. We counted the number of neuronal components with averaged ΔF/F responses that increased by 50% or more compared with before the stimulus, which we refer to as “activated”, and those that decreased by 50% or more, which we refer to as “inhibited” (***Figure 2A, B***). In response to IR heating, the ratio of activated to inhibited neuronal components (A/I) was 5.0 for Hb, 2.0 for Rl, and 4.8 for Vm (**Fig. 2C**). Similarly, responses to electric shock had A/I ratios of 2.8 for Hb, 1.8 for Rl, and 4.0 for Vm (***Figure 2D***).

**Figure 2.**
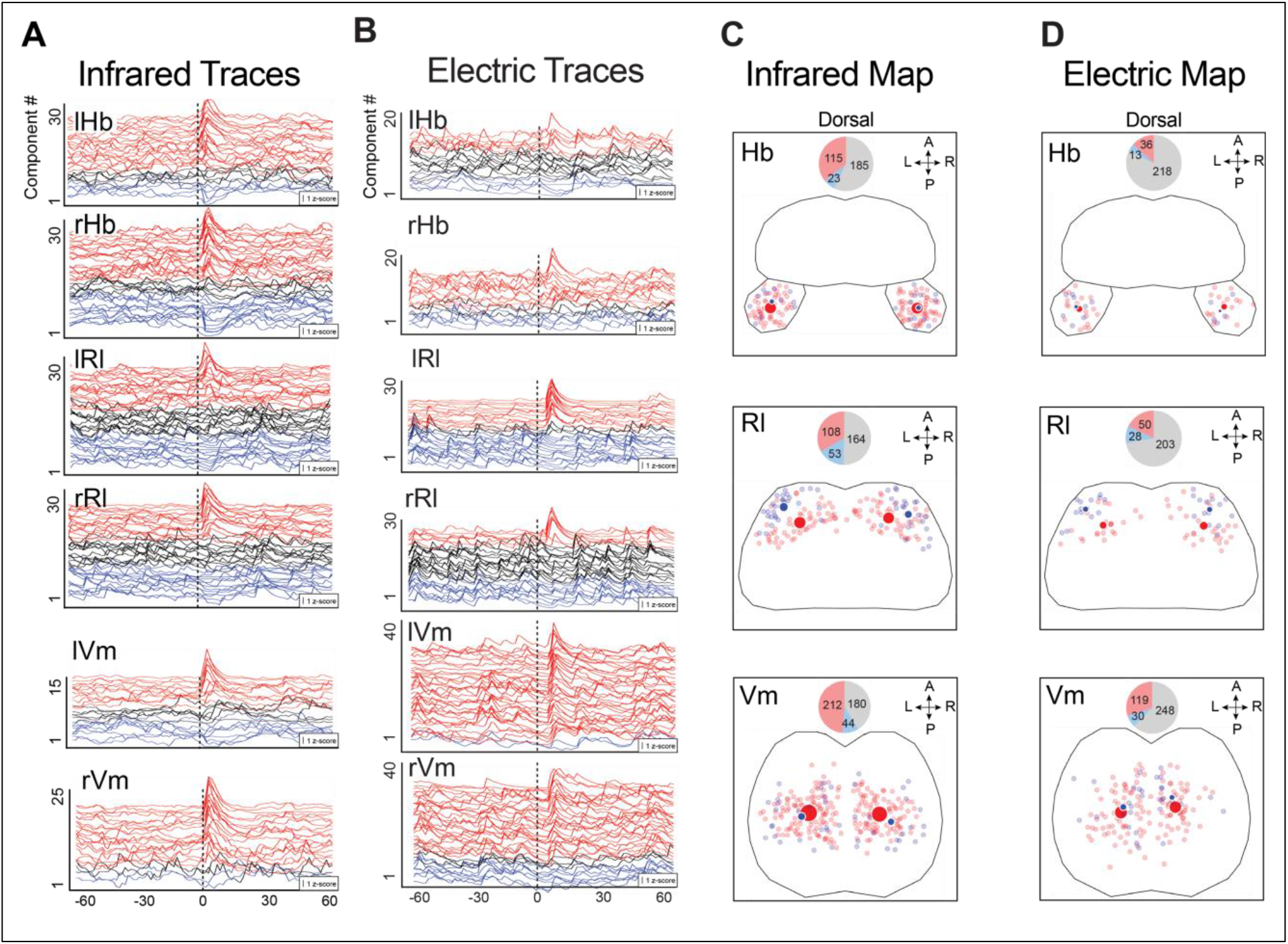
Response of neuronal components to noxious heat stimuli and electric shock. (**A**) Neuronal component z-score (μ/σ, ΔF/F) traces from an example fish as determined by CaImAn reveal a diverse response upon IR stimulation at the vertical dotted line. Red denotes activation (increase in activity ≥ 50% of background ΔF/F); blue, inhibition (decrease in activity ≤ –50% of background ΔF/F); Black, neither. Order is determined by z-score. (**B**) Neuronal component z-score traces as in (A) in response to electric shock at the vertical dotted line. (**C**) Locations of activated (small red dots) and inhibited (small blue dots) neuronal components in response to heating with an IR laser for all fish tested. Large dots are centers of mass of corresponding neuronal components. Size of the large dots is proportional to the number of activated or inhibited neuronal components. Pie graphs (inset) show the number of activated (red) inhibited (blue), and unresponsive (grey) neuronal components. (**D**) Same as (C), but for neurons that respond to electric shock. Abbreviations: A, anterior; AUC, area under the curve; L, left; lHb, left habenula; lRl, left rostrolateral; lVm, left ventromedial; P, posterior; R, right; rHb, right habenula; rRl, right rostrolateral; rVm, right ventromedial. **Figure 2-source data 1**. Numerical source data for graphs in ***Figure 2A*** and ***2B***, and *plots in* ***Figure 2C*** and ***2D***. **Figure 2-figure supplement 1.** Overlay of center of mass for excited and inhibited neuronal components in response to IR and electricity.

Although it was not possible to expose the fish to multiple stimuli within a single imaging session due to their fragility, we plotted the cumulative distribution of activated vs. inhibited neuronal components onto a canonical fish template for both IR heating and electric shock so that we could compare responses to the two stimuli (***Figure 2C, D, Figure 2-figure supplement 1***). In Rl, components activated by both IR heating and electric shock were generally located in the same region, caudal and medial to the region containing inhibited components.

### Distinct responses to full– and partial-looming stimuli in Rl

Given that two different noxious stimuli evoked activity in Hb, Rl, and Vm (***Figure 1, 2***), we asked how the same regions would respond to a non-noxious stimulus that still indicated an imminent threat of bodily harm. For many visual species, including zebrafish, a looming stimulus consisting of a black disc that increases in size, mimicking the shadow of an approaching predator, signals a potential threat and triggers stereotyped escape behaviors (Temizer et al., 2015). To generate a looming stimulus, we projected a black disc onto a small projector screen located to the right of the 7–9 dpf zebrafish, which started as a 1**°** point and grew over 500 ms to cover the fish’s entire visual field. After remaining at full size for 5 s, the disc contracted back to 1**°**. During this sequence, we monitored GCaMP fluorescence. As with exposure to IR heating and electric shock, the full-looming stimulus immediately elicited increased activation in all six regions activated by noxious stimuli (***Figure 3A-D***). There was residual activity in several areas, including rVm, rRl, and lRl 8 s after the stimulus ceased. Quantitation of the average ΔF/F before and after presentations showed significant increases in all three areas (n = 5, ***Figure 3E***).

**Figure 3.**
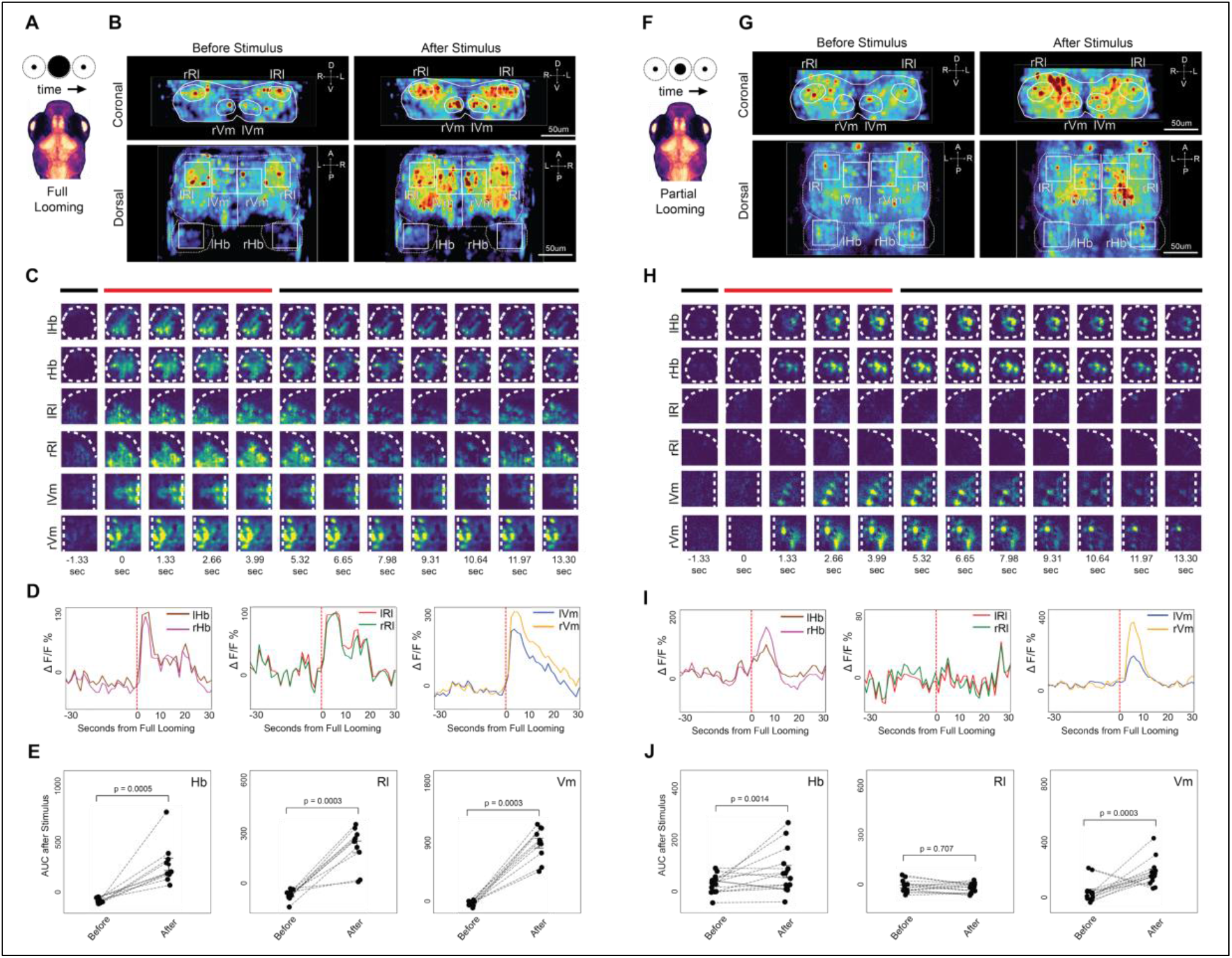
Neuronal activation in response to full– and partial-looming stimuli. (**A**) Schematic of a 7 dpf zebrafish subjected to a full-looming disc stimulus. (**B**) Heatmap of GCaMP fluorescence in a representative zebrafish immediately before and after a full looming stimulus, averaged over 5 presentations, shows an increase in average activity across all brain areas previously found to be activated by noxious stimuli (rHb, lHb, rRl, lRl, rVm, lVm). (**C**) Timelapse of GCaMP fluorescence heatmaps for the six areas outlined in (B) shows an abrupt increase in activity that commences with the full-looming stimulus. (**D**) Region-specific ΔF/F activity traces from a representative fish, averaged across 5 trials, display pronounced increases in all measured areas following IR laser exposure. (**E**) Mean increase above background ± SEM for ΔF/F A.U.C. for 5 frames, 6.65 seconds for each fish for each brain region outlined in (B). (**F**) Schematic of a 7 dpf zebrafish subjected to a partial-looming disc stimulus. (**G**) Heatmap of GCaMP fluorescence in a representative zebrafish immediately before and after a partial-looming stimulus, averaged over 3 presentations. (**H**) Time-lapse of GCaMP fluorescence heatmaps for the six areas outlined in (B) shows an increase in activity in the Hb and Vm regions, but no apparent change in activity in the Rl region. Activity in Hb and Vm was sustained for over 10 seconds. (**I**) Region-specific ΔF/F and activity traces from a representative fish, averaged across 3 trials, display significant activity increases in Hb and Vm, but not in Rl, in response to the partial looming stimulus. (**J**) Mean increase above background ± SEM for ΔF/F after the partial-looming stimulus (AUC; 5 frames, 6.65 seconds). All statistical comparisons made with one-tailed t-test. The red dashed lines denote the onset of the stimulus. Abbreviations: A, anterior; AUC, area under the curve; D, dorsal; L, left; lHb, left habenula; lRl, left rostrolateral; lVm, left ventromedial; P, posterior; R, right; rHb, right habenula; rRl, right rostrolateral; rVm, right ventromedial; V, ventral. **Figure 3-source data 1**. Numerical source data for graphs in ***Figure 3D***, ***3E***, ***3I***, and ***3J***.

Given the robust response to the full-looming stimulus, we asked whether a similar, but less intense stimulus, might elicit a different neuronal response. Thus, we developed a partial looming stimulus, which is similar to the full looming stimulus, except the disc stops growing after 300 ms when it covers approximately 48 degrees of the visual field, and then, after 5 s, reduces in size over 300 ms to a 1**°** point, which could mimic a predator that comes close to the fish but then stops (***Figure 3F***). Strikingly, the partial-looming stimulus did not elicit a significant overall ΔF/F increase in Rl, although increases in activity in Hb and Vm were significant (n = 7, ***Figure 3G-J***).

A/I ratios of individual activated and inhibited neuronal components responding to the full-looming stimulus were 7.2 in Hb, 1.4 in Rl, and 5.5 in Vm, indicating that activation predominated overall (***Figure 4A, C***). However, in response to the partial-looming stimulus, neuronal components showed an A/I of 1.2 in Hb and 3.0 in Vm, whereas inhibited components predominated in Rl (A/I = 0.41; ***Figure 4B, D***). Thus, multiple measures of neuronal activity are consistent with Rl being activated by noxious stimuli and a full-looming stimulus, but not by a partial-looming stimulus. In contrast, Vm and Hb are activated by all stimuli tested.

**Figure 4.**
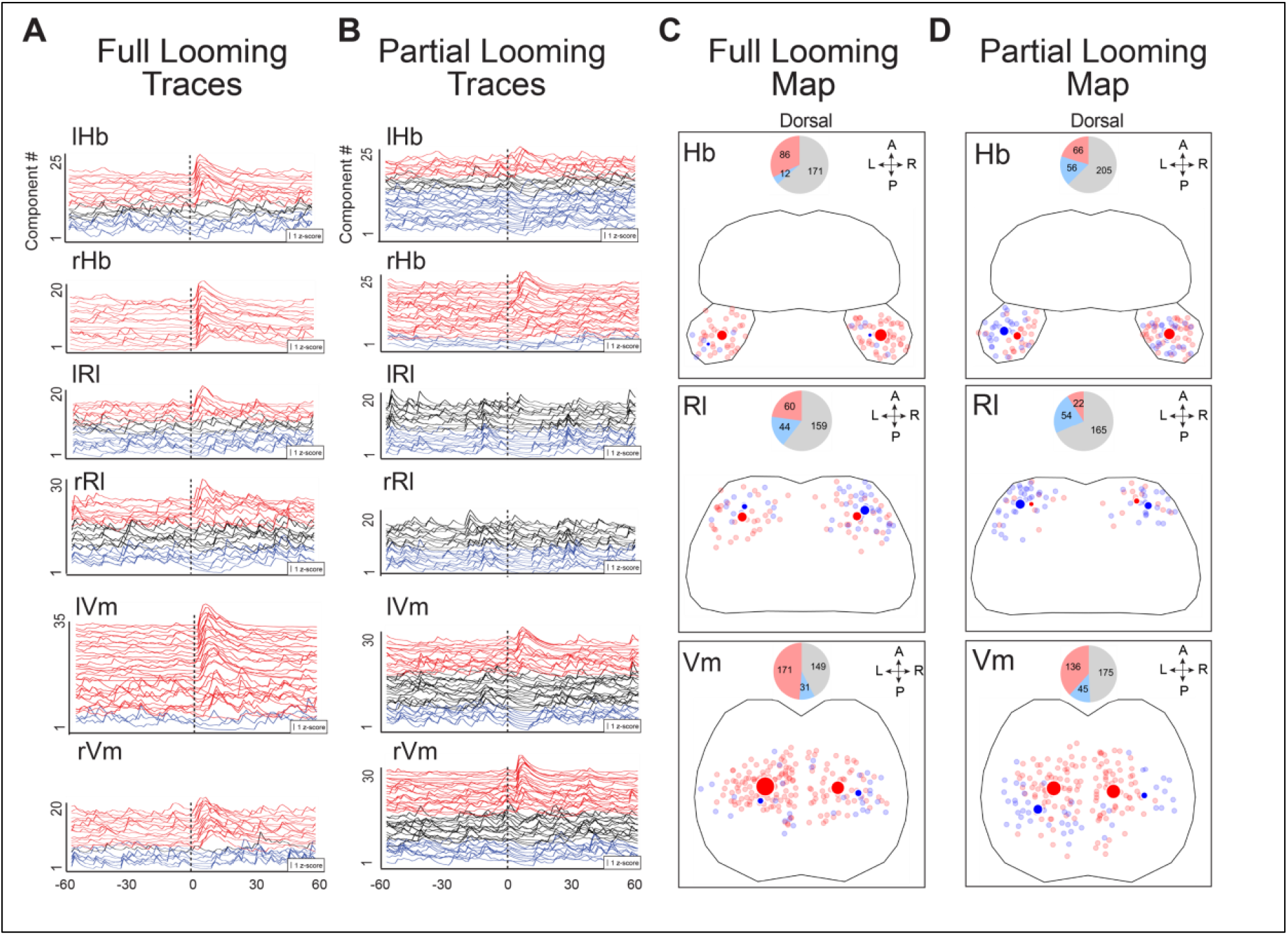
Response of neuronal components to full and partial looming stimuli. (**A**) Neuronal component z-score (μ/σ, ΔF/F) traces for an exemplary fish in response to a full looming stimulus at the vertical dotted line, averaged over 5 presentations. Red denotes activation (increase in activity ≥ 50% of background ΔF/F); blue, inhibition (decrease in activity ≤ –50% of background ΔF/F); black, neither. (**B**) Neuronal component z-score as in (A) in response to a partial looming stimulus averaged over 3 presentations in an exemplary fish. Activated and inhibited traces are seen in Hb and Vm, but not in Rl. (**C**) Locations of neurons that are activated (small red dots) and inhibited (small blue dots) in response to a full looming stimulus for all fish tested. Large dots are centers of mass of corresponding neuronal components. Size of the large dots is proportional to the number of activated or inhibited neuronal components. Pie graphs (inset) show the number of activated (red), inhibited (blue), and unresponsive (grey) neuronal components. (**D**) Same as (C) in response to a partial looming stimulus for all fish tested. Abbreviations: A, anterior; AUC, area under the curve; L, left; lHb, left habenula; lRl, left rostrolateral; lVm, left ventromedial; P, posterior; R, right; rHb, right habenula; rRl, right rostrolateral; rVm, right ventromedial. **Figure 4-source data 1.** Numerical source data for graphs in ***Figure 4A*** and ***4B***, and plots in ***Figure 4C*** and ***4D***.

### Mild negative-valence stimuli do not activate Rl neurons

To further test whether Rl neurons respond only to highly threatening stimuli, such as noxious stimuli, or to a full-looming stimulus, we exposed larval zebrafish to stimuli that could be perceived as threatening but do not indicate imminent bodily harm and measured the response in Hb, Rl, and Vm. Specifically, we exposed the zebrafish to sudden vibration, loud noises, and a transition from light to darkness. Each of these stimuli has been shown to elicit tail movements or increases in turning angle in zebrafish (Emran et al., 2008; Neo et al., 2015; Ro et al., 2022), consistent with their being inherently threatening. To expose the fish to vibration while recording GCaMP activity with the SPIM, we tapped the right side of the imaging chamber with a plastic stick for 5 s. We also presented an auditory stimulus by sounding an airhorn that produced a 120-130 dB sound lasting 0.5 s. Finally, fish that had been equilibrated to darkness, then exposed to light for 4 s, followed by an abrupt return to darkness, were analyzed for response to the light-to-dark transition. Each stimulus was repeated 5 times with a 2-min interval between trials. During and after each stimulus, increased activity was apparent in Hb and Vm, whereas no overall activity change was observed in Rl (***Figure 5A-I***).

**Figure 5.**
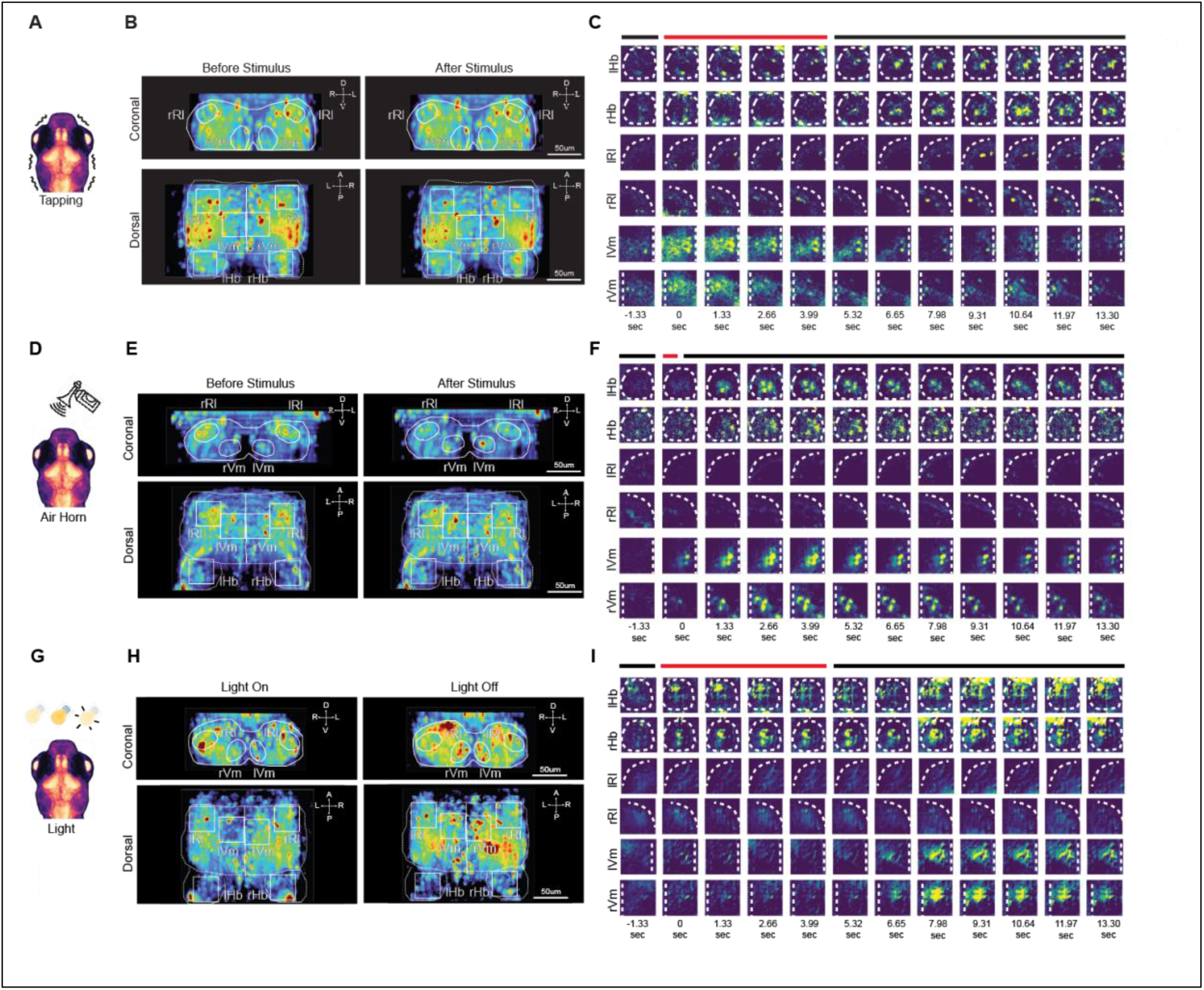
Neuronal activation in response to vibration, a loud noise, and light. (**A**) Schematic of a 7 dpf zebrafish subjected to vibration. (**B**) Heatmap of GCaMP fluorescence in a representative zebrafish immediately before and after a vibration stimulus, averaged over 5 presentations, shows an increase in average activity across Hb and Vm, but not Rl. (**C**) Timelapse of GCaMP fluorescence heatmaps for areas outlined in (B) show an increase in activity that commences with the initiation of tapping and grows until 14 s after cessation of tapping in Hb. In Vm the activity increases during the tapping stimulus and decreases abruptly after cessation. There is no apparent increase in activity in Rl. (**D**) Schematic of a 7 dpf zebrafish subjected to the blast of an air horn. (**E**) Heatmap of GCaMP fluorescence in a representative zebrafish immediately before and after an air horn blast, averaged over 5 presentations, shows an increase in average activity across Hb and Vm, but not Rl. (**F**) Timelapse of GCaMP fluorescence heatmaps for areas outlined in (E) show an increase in activity that commences with the onset of the loud noise in Hb and Vm and lasts for over 10 s. There is no apparent increase in activity in Rl. (**G**) Schematic of a 7 dpf zebrafish that is first equilibrated to darkness, then exposed to light for 4 s and then returned to darkness. (**H**) Heatmap of GCaMP fluorescence in a representative zebrafish immediately before and after transitioning from light to darkness, averaged over 5 presentations shows an increase in average activity across Hb and Vm, but not Rl. (I) Timelapse of GCaMP fluorescence heatmaps for areas outlined in (H) shows an increase in activity in Hb and Vm that commences ∼ 2 s after the onset of darkness and lasts for over 5 s. There is no apparent increase in activity in Rl. Abbreviations: A, anterior; D, dorsal; L, left; lHb, left habenula; lRl, left rostrolateral; lVm, left ventromedial; P, posterior; R, right; rHb, right habenula; rRl, right rostrolateral; rVm, right ventromedial; V, ventral.

Statistical analysis showed that overall ΔF/F increased significantly in the Vm in response to tapping, but activity in Rl and Hb did not increase significantly (n = 6 fish, ***Figure 6A, B***). Similar results were found for exposure to the air horn blast where activity significantly increased in Hb and Vm, but not in Rl (n = 4 fish, ***Figure 6C, D***). Finally, overall ΔF/F increased significantly in Hb and VM, but not in Rl following a light to dark transition (n = 5, ***Figure 6E, F***). Consistent with these results, A/I ratios for the neuronal components found in Hb and Vm ranged from 1.6-4.3 but were substantially less than 1 (0.29-0.66) in Rl for tapping, airhorn, and light-to-dark transitions (***Figure 7A-F***). Thus, overall activity in Hb and Vm increases significantly in response to moderately threatening stimuli (except for Hb in response to tapping) and activated neuronal components outnumber those that are inhibited. In contrast, in Rl, overall ΔF/F does not increase significantly, and inhibited neuronal components outnumber activated ones.

**Figure 6.**
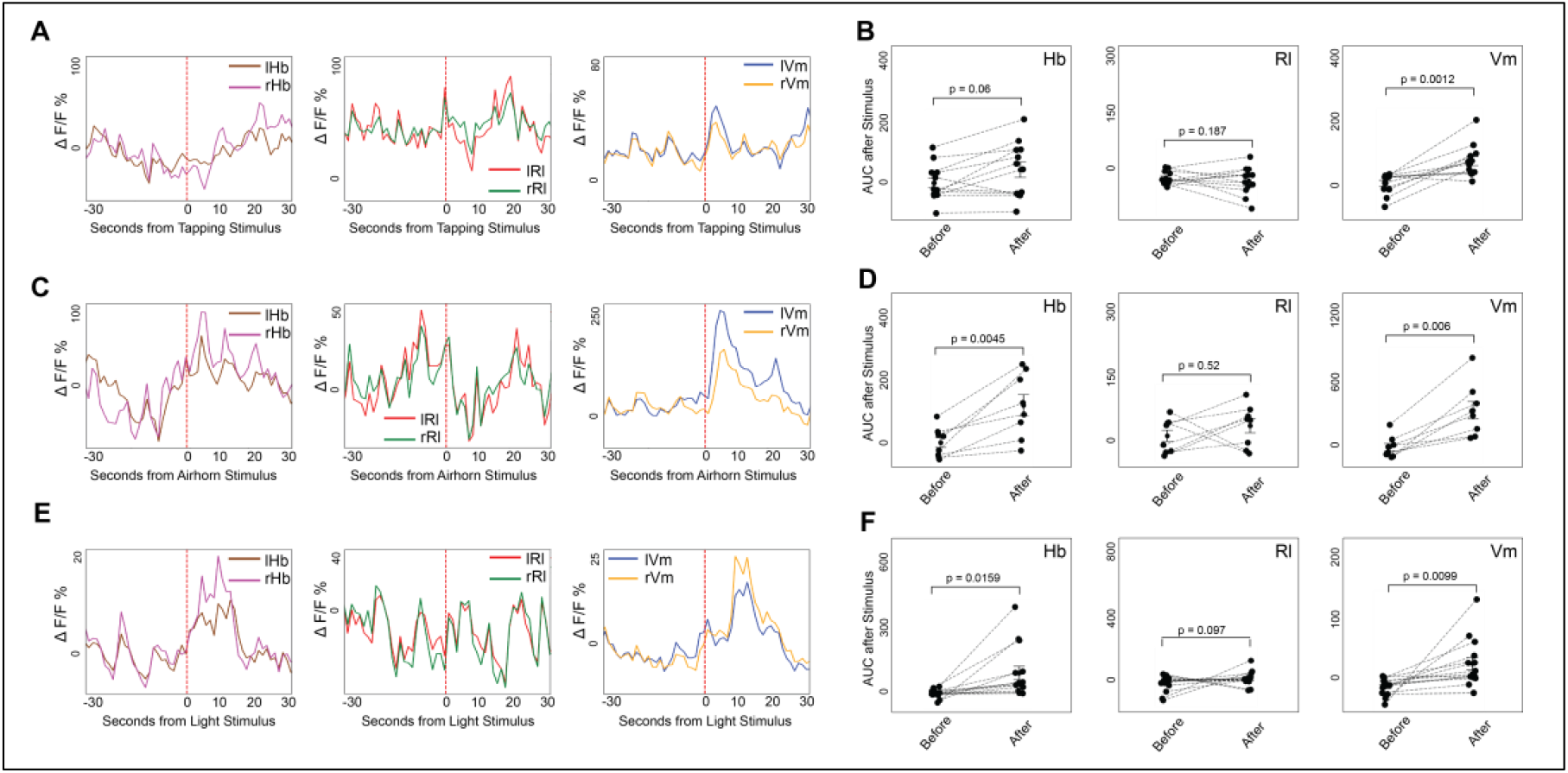
Response of total ΔF/F within different regions to vibration, a loud noise, and light. (**A**) Average ΔF/F activity traces within specific regions from a representative fish, averaged across 5 trials in response to a tapping stimulus. (**B**) Activity rates for 6 fish before and after the tapping stimulus (AUC; 5 frames, 6.65 seconds). Significant increases in activity were seen in Vm but not in Hb and Rl. (**C**) Average ΔF/F activity traces within specific regions from a representative fish, averaged across 5 trials in response to a blast from an airhorn. (**D**) Responses to a blast from an airhorn from 4 fish (AUC; 5 frames, 6.65 seconds). Significant increases in activity were seen in HB and Vm, but not in Rl. (**E**) Average ΔF/F activity traces within specific regions from a representative fish, averaged across 5 trials in response to a light-to-dark transition. (**F**) Responses to a light-to-dark transition from 6 fish (AUC; 7 frames, 9.31 seconds). Significant increases in activity were seen in HB and Vm, but not in Rl. The red dashed lines denote the onset of the stimulus. Abbreviations: AUC, area under the curve; lHb, left habenula; lRl, left rostrolateral; lVm, left ventromedial; P, posterior; rHb, right habenula; rRl, right rostrolateral; rVm, right ventromedial. **Figure 6-source data 1.** Numerical source data for graphs in ***Figure 6A-F***.

**Figure 7.**
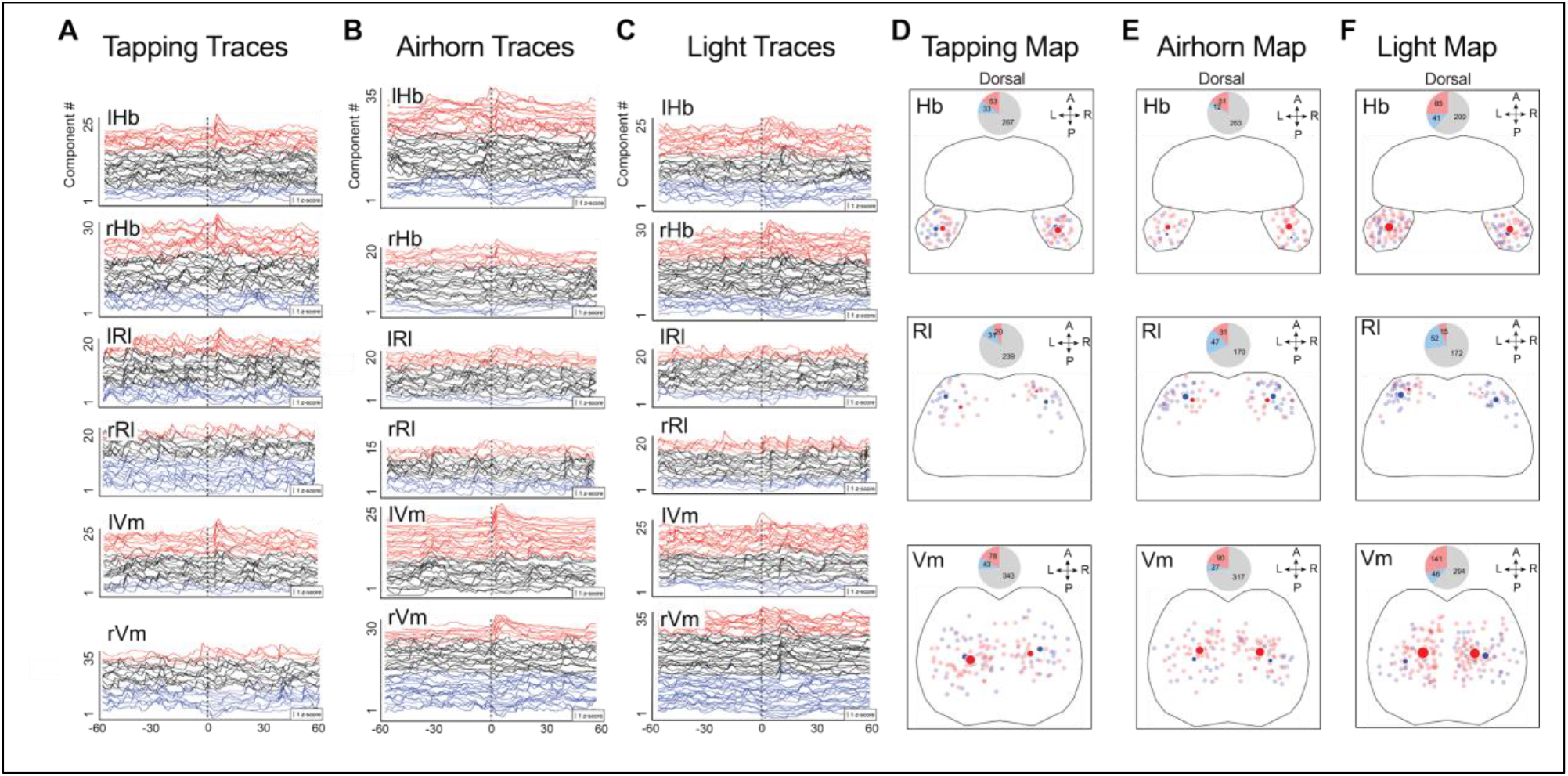
Response of neuronal components to vibration, a loud noise, and light. (**A**) Neuronal component z-score (μ/σ, ΔF/F) traces of a representative fish in response to tapping at the vertical dotted line. Red denotes activation (increase in activity ≥ 50% of background ΔF/F); blue, inhibition (decrease in activity ≤ –50% of background ΔF/F); black, neither. Order is determined by z-score (μ/σ), which measures the magnitude and consistency of the response. In Hb and Vm, the responses of neuronal components to all three stimuli tend to be better defined than those in Rl. (**B, C**) Neuronal component z-score traces as in (A), but in response to the blast of an airhorn (B) or a transition from light to dark (C) at the vertical dotted line. (**D-F**) Locations of neuronal components in Hb, Rl, and Vm. Small puncta mark locations where neuronal components respond by increasing (red) or decreasing (blue) activity in response to tapping (D), a blast from an airhorn (E) or a transition from light to dark (F). Large dots represent centers of mass of corresponding neuronal components. Size of the large dots represents the number of activated or inhibited neuronal components. Pie graphs (inset) show the number of activated (red), inhibited (blue), and unresponsive (grey) neuronal components. Note that in Rl, inhibited neurons outnumber activated neurons, whereas in Hb and Vm, the proportion of activated components are roughly equal to or greater than that of inhibited components. Abbreviations: A, anterior; L, left; lHb, left habenula; lRl, left rostrolateral; lVm, left ventromedial; P, posterior; R, right; rHb, right habenula; rRl, right rostrolateral; rVm, right ventromedial. **Figure 7_source data 1.** Numerical source data for graphs in ***Figure 7A-C***, and plots in Figure 7C-E

In conclusion, our results suggest that Rl responds exclusively to the most threatening stimuli, but not to less threatening stimuli both as indicated by overall activity and A/I ratio of neuronal components (***Figure 8***). In contrast, Hb and Vm respond to all negative-valence stimuli and neuronal components have A/I ratios > 1 (***Figure 8***), regardless of whether they are strongly or weakly threatening. Thus, in principle, signals present in Rl could be used to identify highly threatening stimuli. Tail movements were imaged before and after presentation of the stimuli. It was found that electricity, tapping, and a loud noise emitted by an airhorn all caused a significant increase in twitches (***Figure 8-figure supplement 1***). Surprisingly, rates of twitching did not change before vs. after the other stimuli (IR heating, full looming, partial looming, and dark-to-light transitions).

**Figure 8.**
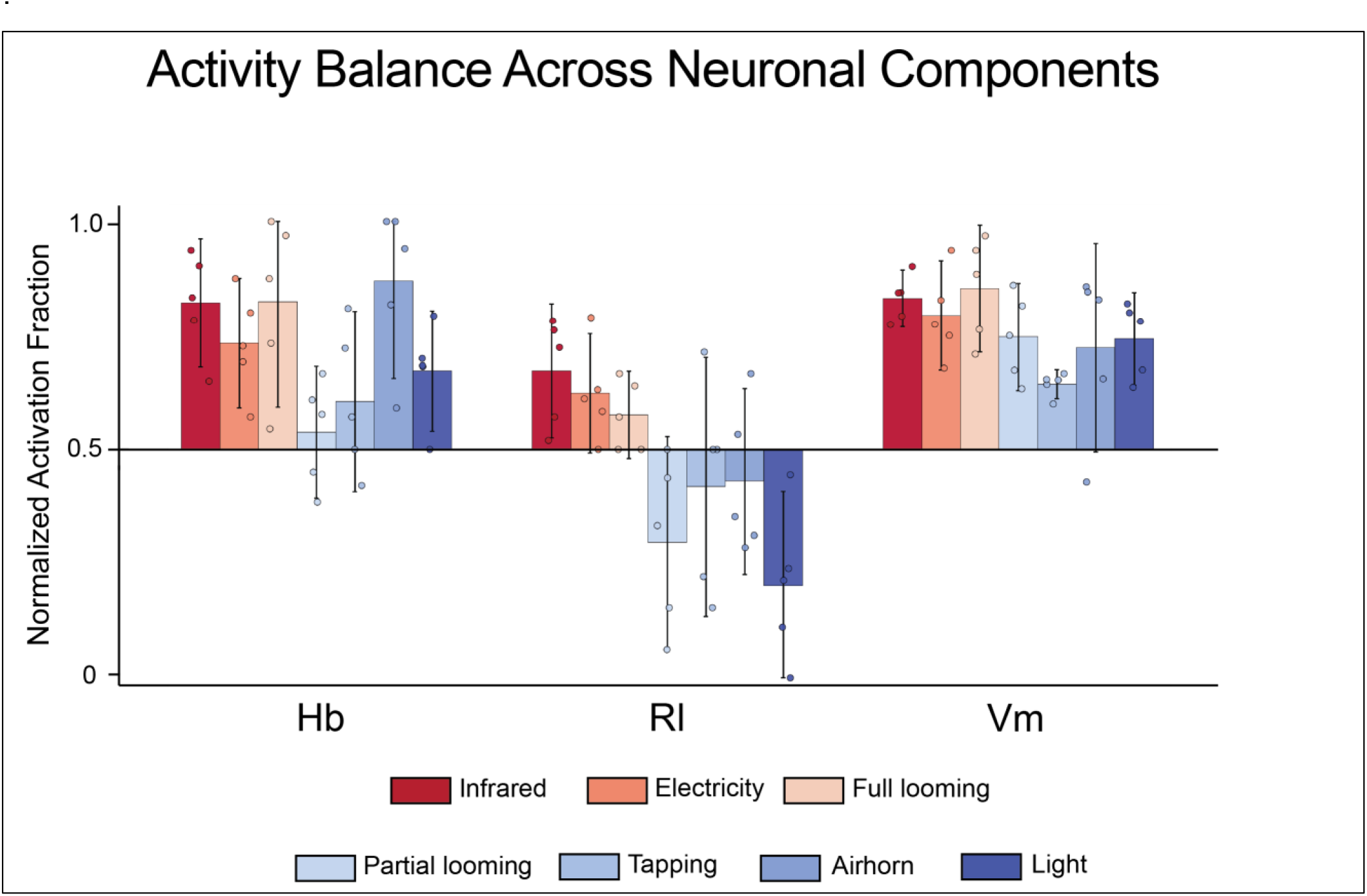
Rl is activated by highly threatening stimuli but not by mildly threatening stimuli. The ratio of activated to inhibited neuronal components (A/I) in Rl is greater than 1 for highly threatening stimuli but less than 1 for mildly aversive stimuli. In contrast, in Hb and Vm, the A/I ration is greater than one for all aversive stimuli. Dots represent A/I for individual fish (biological replicates). Bars represent 95% confidence intervals. Abbreviations: Hb, habenula; Rl, rostrolateral; Vm, ventromedial. **Figure 8-figure supplement 1** Twitches in Response to Stimuli. Number of twitches of individual fish before and after stimuli. Rates (probability) of twitching before and after were compared with a one-tailed t-test. *, p < 0.05; **, p < 0.01; ns, p > 0.05. **Figure 8 Source Data** Numerical source data for graph in **Figure 8**.

### Tiam2a-positive neurons in the rostrolateral pallium

To better understand the properties of neurons in the Rl area that respond specifically to imminent threats, we used the MapZebrain cellular-resolution atlas of the larval zebrafish brain (Kunst et al., 2019). We found that in 6 dpf transgenic fish (y264Et, see methods), the *tiam2a* promoter drives GFP specifically in a region of the pallium that matches the location of Rl neurons. *Tiam2a* is the zebrafish homolog of the mammalian gene *Tiam2*, which encodes a Rac1-associated GEF (Chiu et al., 1999).

To test whether *tiam2a* expression is a marker for neurons that respond to threatening stimuli, we exposed 7–9 dpf Tg[tiam2a:GFP] larval zebrafish to heating with an IR laser. Then we performed wholemount immunocytochemistry to detect pERK, a marker of neuronal activity. A projection image of GFP and pERK shows striking colocalization between the two markers, with virtually every GFP+ neuron expressing pERK (***Figure 9A, B***). In contrast, under control conditions with no noxious stimuli present, we observed little pERK expression in GFP+ neurons (***Figure 9C, D***). To quantify the amount of co-expression between the two, we measured the total intensity of pERK labeling in cells that expressed GFP and subtracted the background intensity of pERK labeling measured in cells in a nearby region of the fish brain that did not express *tiam2a*. pERK labeling in GFP+ cells was ∼20-fold higher in the IR-laser-exposed group (n = 6) than in the control group (n = 8), a statistically significant difference (one-tailed t-test; ***Figure 9E***). These data are consistent with *tiam2a* being a marker for pallial neurons that are more active when zebrafish are exposed to noxious stimuli.

**Figure 9.**
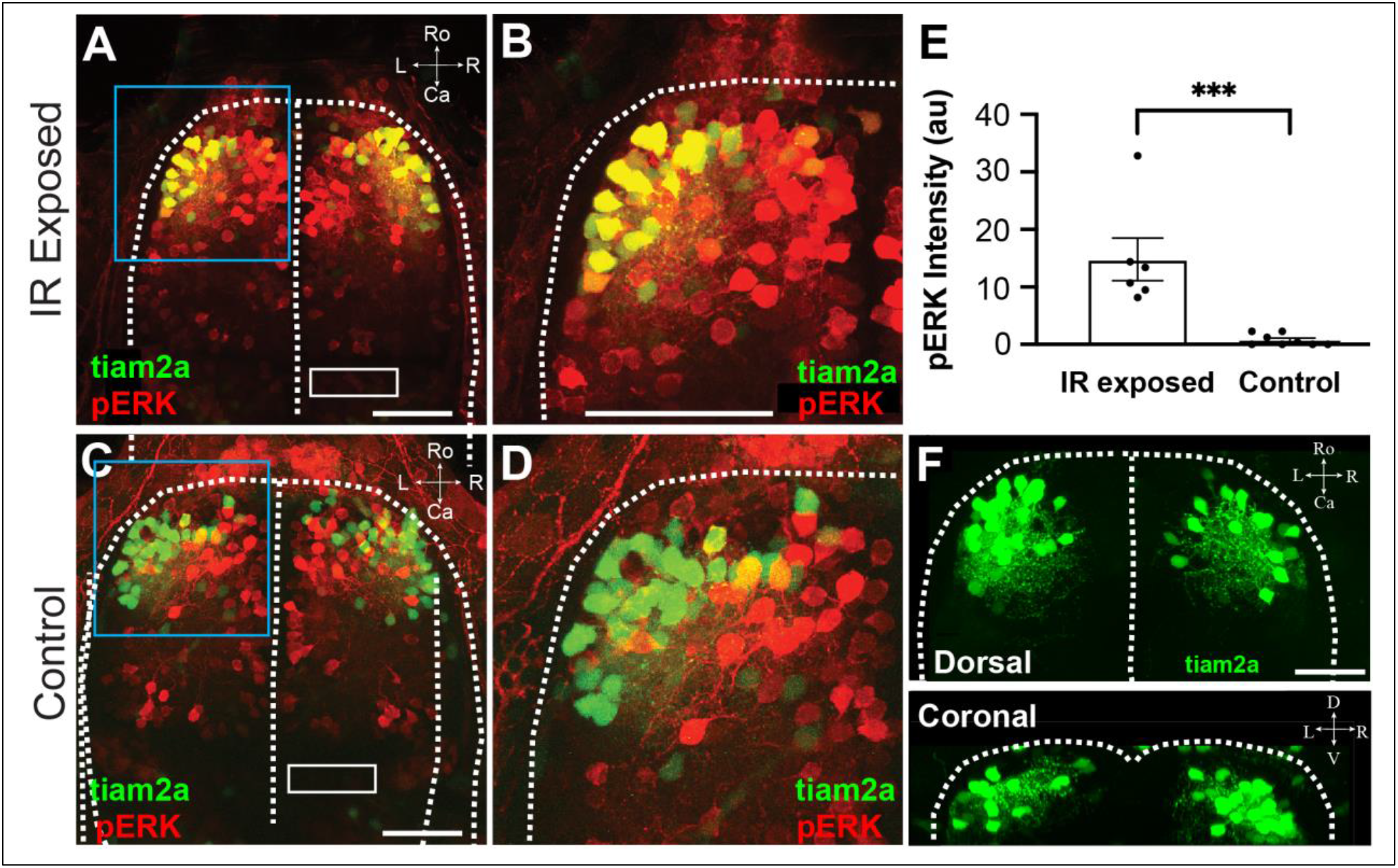
Cells expressing tiam2a become active when exposed to IR heating. (**A**) 7 dpf Tg[tiam2a:GFP] larval zebrafish exposed 10 times to heating with an IR laser and then immunostained for pERK to label active neurons. Virtually all GFP-positive neurons were co-labeled with pERK, suggesting that Tiam2a-positive cells are activated by a painful stimulus. (**B**) Close-up of the area bordered by the blue rectangle in (A). (**C**) 7 dpf Tg[tiam2a:GFP] larval zebrafish control that was not exposed to heating showed very little overlap between GFP labeling and labeling for pERK. (**D**) Closeup of the area bordered by the blue rectangle in (C). (**E**) Average intensity of pERK immunostaining in Tiam2a+ cells, with background (measured in the white rectangle in (A)) subtracted, for fish exposed to IR heating (n = 6) and control fish (n = 8). Significantly more labeling was found in fish exposed to heating (P = 0.0009, t-test) vs. in control fish. (**F**) Tg[tiam2a:GFP] fish showing cell bodies concentrated in the rostrolateral area and local axonal connections. Scale bar 50 μm. Abbreviations: au, arbitrary units; Ca, caudal; D, dorsal; IR, infrared; L, left; R, right; Ro, rostral. **Figure 9_source data 1**. Numerical source data for graph in **Figure 9E**.

To investigate the morphology of Rl neurons, we examined Tg[tiam2a:GFP] fish (***Figure 9F***). From the dorsal view, GFP-labeled neurons are highly clustered in the rostrolateral pallium, and from the coronal view, they appear superficial and dorsal. Furthermore, they have short axons, suggesting they are involved in local connectivity. Thus, *tiam2a* promoter-mediated expression defines a tightly packed group of neurons that respond to a noxious stimulus in the 7–9 dpf larval zebrafish. These neurons are likely negative-valence neurons that respond specifically to a variety of highly threatening stimuli, but not to less threatening aversive stimuli.

## Discussion

In this study, we defined responses of neurons in the larval zebrafish forebrain to noxious stimuli (IR heating and electric shock), non-noxious threatening stimuli (full looming), and less threatening stimuli (partial looming, vibration, loud noise, and light-to-dark transitions). Prior research has shown that in the adult zebrafish, neurons in the lateral zone of the dorsal telencephalic area, which partially overlaps with Rl, drive ongoing habenular activity (Bartoszek et al., 2021). However, dynamic responses of forebrain neurons to negative-valence stimuli have not been reported previously. Here, we have found that noxious and non-noxious threatening stimuli activate a distinct population of tightly clustered neurons within the rostrolateral dorsal pallium. However, neurons within this same region do not respond to less threatening stimuli. In contrast, neurons in the ventromedial pallium and the habenula respond to all the stimuli tested, encompassing both highly threatening and mildly aversive stimuli.

In addition to neuronal components activated by negative-valence stimuli, we found components in Rl that exhibited reduced activity upon exposure to noxious and threatening stimuli. Previously, it was found that neurons inhibited by negative-valence stimuli may also be activated by positive-valence stimuli, suggesting that these neurons may respond to positive valence (Belova et al., 2008; Smith and Torregrossa, 2021). The ratio of activated to inhibited neuronal components was highly correlated with whether the overall signal from a given region increased in response to a specific stimulus. A/I ratio (number of activated vs. inhibited neuronal components) > 1 was correlated with regional activation, whereas A/I < 1 was associated with a lack of activation. In the future, it will be important to test positive-valence stimuli to determine whether they elicit responses in neurons inhibited by negative-valence stimuli. Attempts in this study to elicit a response from forebrain neurons to the smell of food, a positive-valence stimulus, were unsuccessful (data not shown). Additionally, it will be important to determine whether individual neurons in Vm and Hb are broadly tuned to negative-valence stimuli or respond to stimuli of all valences.

For cells to be classified as negative-valence, they must respond to negative-valence stimuli of all types (Beyeler et al., 2018). We found that a full-coverage looming shadow activated Rl neurons, along with ventromedial and habenular regions, in a manner similar to physically noxious stimuli. However, a partial looming shadow failed to elicit a response in Rl, although it activated Hb and Vm. Surprisingly, a previous study that examined looming stimuli in zebrafish did not find a significant number of pallial neurons that responded to looming stimuli (Fotowat and Engert, 2023) in contrast to the results presented here. This apparent discrepancy may be accounted for by differences between the two studies in looming stimuli presented, transgenic GCaMP-expressing fish, and in the type of microscope used for imaging.

Since Rl responds to negative-valence stimuli that in some contexts can evoke tail twitches, it is possible that Rl provides input to hindbrain escape circuits (Kunst et al., 2019). Application of DiI (a lipophilic fluorescent tracer) crystals in the dorsal/dorsolateral pallium, where Rl is located, resulted in labeled fibers within the lateral forebrain bundle (Yáñez et al., 2022). The lateral forebrain tract projects to the diencephalon, but not the hindbrain (Rink and Wullimann, 2004), implying that any connections from Rl to the hindbrain are likely indirect. This is consistent with our findings that there is not a one-to-one correlation between activation of Rl and increase in the rate of twitching. However, delineating a pathway from Rl to brainstem escape circuits, even if indirect, would be important for understanding how Rl contributes to escape behavior.

In this study, we exploited the unique properties of the larval zebrafish to obtain parallel recordings of neurons throughout the forebrain in response to negative-valence stimuli. One drawback of using larval zebrafish is that they are relatively delicate, making it difficult to present multiple stimuli to the same fish. Consequently, although it is possible to conclude that neurons in Rl are likely to respond when the fish is exposed to one of several different stimuli, we cannot conclude that individual neurons respond to more than one stimulus. Although tail twitches and heart rate were monitored with every stimulus, we were unable to detect any significant changes in heart rate (data not shown). Electric shock, vibration, and a loud noise elicited a significant increase in the probability of twitching consistent with previous results (***Figure 8-figure supplement 1***; Kenney et al., 2017; Manuel et al., 2014; Neo et al., 2015; Ro et al., 2022). However, we did not elicit an increase in tail twitching in response to looming stimuli, IR heating, and light-to-dark transitions, although these stimuli elicited responses in other contexts (Dempsey et al., 2022; Emran et al., 2008; Temizer et al., 2015). One possible explanation for these results is that when the zebrafish is mounted on the SPIM with its head encased in agarose while being illuminated by a 2P light sheet, it no longer responds to these stimuli as it does in other contexts. A goal for future work is to develop imaging systems that can non-invasively record activity throughout the zebrafish brain in real time while still allowing the fish to remain in a natural environment. Remarkably, forebrain activity, particularly in Rl, appears to correlate well with the apparent severity of the stimuli, suggesting that it might be a better indicator of the internal state of the zebrafish’s brain than behavior. Together, our results suggest that a transgenic zebrafish expressing GCaMP under the *tiam2a* promoter could provide real-time insight into the animal’s internal state.

Several studies have suggested that the dorsomedial pallium in teleost fish, including zebrafish, is the homolog of the amygdala and the dorsolateral pallium is homologous to the hippocampus based on developmental, topological, and molecular arguments (Baker and Wong, 2021; Mueller et al., 2011; von Trotha et al., 2014). In addition, the medial pallium in general and a specific type of pallial neuron were found to be necessary for fear conditioning in zebrafish (Broglio et al., 2005; Lal et al., 2018; Manuel Portavella et al., 2004; Portavella et al., 2002). In contrast, the results from this study and our previous study suggest that Rl neurons are similar in some respects to neurons in the mammalian amygdala. For instance, both of our studies show that Rl neurons respond to negative-valence stimuli, a response also seen in many neurons in the mammalian amygdala (Beyeler et al., 2018). Rl neurons also become responsive to neutral stimuli following a novel form of classical conditioning (Dempsey et al, 2022). Similarly, following fear conditioning, neurons in the mammalian amygdala become responsive to conditioned stimuli that were paired with unconditioned stimuli (Grewe et al., 2017; Zhang and Li, 2018). Finally, neurons within Rl exhibit an increase in excitatory synapses following classical conditioning, and amygdalar neurons undergo long-term potentiation, consistent with synaptic change, following fear conditioning (Dempsey et al., 2022; Rogan et al., 1997). However, unlike the amygdala, where negative– and positive-valence neurons are distributed in a salt and pepper fashion (Beyeler et al., 2018), the negative-valence cells that we found in Rl tend to be tightly clustered (***Figure 9***). Furthermore, *tiam2a* is a homolog of mammalian *Tiam2*, which is highly expressed in the hippocampus, and is less abundant in the amygdala in the mouse (Chiu et al., 1999). Thus, our data suggest that Rl shares attributes of both the amygdala and the hippocampus, and that teleost fish may have unique brain structures with no exact homologs in mammals.

## Author contributions

C.D.S. and D.B.A. designed the experiments and wrote the paper; C.D.S. performed the live-imaging research, contributed new analytic tools, and analyzed data; Z.D. performed the pERK experiments; T.V.T. and S.E.F. supervised the development and maintenance of the SPIM used for live imaging. W.P.D. performed preliminary experiments that were critical for the development of the experimental protocols used in this paper. D.B.A. secured funding and supervised all aspects of the project.

## Materials and Methods

### Zebrafish husbandry and embryo/larval care

Many of the methods presented in this paper were developed previously (Dempsey et al., 2022). Larval and adult Casper mutant zebrafish were maintained in an in-house veterinary facility in accordance with protocols approved by the University of Southern California (USC) Institutional Animal Care and Use Committee (IACUC). All experiments described in this work were approved by the USC IACUC. Embryos were kept in egg water (1.19 g NaCl, 0.377 g CaSO4·2H2O, 265 mL methylene blue, filled up to 5 liters with ddH_2_O) at 28 °C in an incubator with a 13 hr:11 hr light:dark cycle until 5-6 dpf, when they were transferred to tanks within the husbandry facility. On the husbandry system, zebrafish were also maintained on a 13-h dark/11-h light cycle. The air and water temperature in the zebrafish facility are maintained at 26-28°C. Zebrafish larvae used in experiments (7–9 dpf) were transferred between the facility and the experimental apparatus in zebrafish husbandry system water (HSW) and kept at 26°C – 28°C between experiments.

### Tg[CAG_NRSE_:GCaMP6s]

To monitor cell Ca^++^ activity in these experiments, we used the transgenic zebrafish Tg[CAG_NRSE_:GCaMP6s] with GCaMP6s (Chen et al., 2013) expression driven by a β-Actin promoter (Quitschke et al., 1989) to drive widespread expression and 3X NRSE (neuron-restrictive silencer element) to restrict expression to neurons (Bergeron et al., 2012).

### Tg[tiam2a:GFP]

The transgenic fish y264Et (originally created by the lab of Harold Burgess), with the construct Et(SCP1:GAL4FF), uniquely labels Tiam2a-positive cells with GFP within the rostrolateral pallium (Sprague et al., 2003).

This transgenic fish was deposited in the ZFIN database (Ruzicka et al., 2015) with the following citation: Marquart, G.D., Tabor, K.M., Brown, M., Strykowski, J.L., Varshney, G.K., LaFave, M.C., Mueller, T., Burgess, S.H., Higashijima, S., and Burgess, H.A. (2015) Enhancer Trap Transgenics. ZFIN Direct Data Submission. (http://zfin.org).

### Mounting Zebrafish in Caddie for Imaging on FlexSPIM

To prepare for behavioral experiments, zebrafish subjects were first embedded in a custom behavioral training caddie generated by Protolabs (Stereolithography, ABS-like Translucent Clear [WaterShed]). Specs: ABS-Like Translucent Clear (WaterShed), Normal Res, Natural, Stereolithography, X: 0.980in Y: 0.206in Z: 0.311in. The caddie was filled with 200 mL of 1% (w/v) agarose, melting point ≤ 65°C (Millipore Sigma; SeaPlaque, Cat:50101) dissolved in egg water containing methylene blue, which suppresses fungal growth. During agarose embedding and fluorescence imaging experiments, zebrafish were anesthetized in a final concentration of 70 ppm MS-222 (Western Chemical, Fluka analytical, Cat: A5040-100G) + 70 ppm Isoflurane (Phoenix SKU 012ABB-250CC), an IACUC-approved combination that has been shown to enable fast recovery from anesthesia as compared to using MS-222 alone (Huang WC, et al., 2010). The anesthetics are dissolved in filter sterilized HSW. For all experiments involving agarose-embedding of the zebrafish, a 1.5% (w/v) solution of low melting point agarose (Sea Plaque Agarose, Lonza) dissolved in filter sterilized husbandry system water consisting of instant ocean (ASIN: B000255NKA) and baking soda dissolved in reverse osmosis water to achieve pH 7.2 and a conductance of ∼700 μS) with anesthetic is used. There was no bias in zebrafish sex for larval experiments, as sex determination does not occur in zebrafish before the juvenile stage, ∼24 dpf (Uchida et al., 2002).

As the agarose solidified (before it completely solidified at 5 min), a hair stem (human hair attached to a plastic rod) was used to position the zebrafish so that the body axis was parallel to the surface of the chamber (normal swimming orientation). Once the agarose solidified, the caddie was placed into a 60 mm culture dish (Corning, Cat.: 430166) and 10 ml of fresh filter-sterilized HSW was added. The agarose was then shaved down with a razor such that only the head and body were encased. Before the anesthetic wore off after the introduction of the fresh HSW solution (∼30 seconds), the tail of the zebrafish (just posterior to the swim bladder) was carefully freed from the agarose using fine forceps (biology tip profile, Fine Science Tools, Cat: 11251-10). The rostral aspect of the zebrafish (anterior to the swim bladder) was held firmly in place by the agarose. The caddie was then transferred to another 60 mm culture dish in preparation for transportation to the imaging room.

### FlexSPIM Light Sheet Microscope

After the zebrafish was mounted into the custom Protolabs caddie, it was ready to be placed onto the flexSPIM. The flexSPIM is a versatile, multi-laser twin-microscope system for light-sheet imaging designed and built for multiphoton and visible wavelength fluorescence light sheet excitation (Keomanee-Dizon et al., 2020). The caddie was securely fastened on the dive bar and placed into the chamber filled with 80 ml of fresh, filter-sterilized HSW. During imaging, the objective position was programmed to follow a saw waveform such that the objective took a unidirectional path starting at the top of the fish pallium (0 mm) and transitioned through to the most ventral aspect of pallium (60 µm) at a frequency of 0.75 Hz. The brain camera followed a square waveform that instructs it to acquire 11 images per objective cycle.

### Arduino

For the behavioral experiments, an Arduino unit (Arduino UNO Rev3, Arduino.cc) and the software (Arduino software [IDE]) from the same company were used to control the stimuli. For the IR experiments, a simple program was designed to control the IR laser (NIR RLCO-980-1000-F laser, Roithner Lasertechnik) by turning it on and off for two seconds, every two minutes. For the electricity experiments, a simple program was designed to allow the native Arduino 5V output to be turned on and off for 400 ms every two minutes.

### Volumetric Visualization of Neural Activation

To visualize stimulus-evoked neural activation, volumetric calcium-imaging data were reconstructed into three-dimensional heat maps. Eleven consecutive image planes acquired with the flexSPIM immediately before stimulus onset were combined to generate a baseline 3D fluorescence volume. The same procedure was repeated for frames acquired after the stimulus (the specific time selected was within a 3s post-stimulus window to capture the peak GCaMP response). The pre– and post-stimulus volumes were visualized side-by-side to illustrate regional activation patterns. Each 3D volume was rendered in coronal and dorsal orientations and manually curated to exclude superficial skin layers and non-neuronal regions, providing volumetric heat maps of GCaMP fluorescence before and after stimulation

### Time-Lapse Analysis of Whole Regions

To quantify the temporal dynamics of stimulus-evoked activity within a region as a whole, time-lapse calcium-imaging data were analyzed from 90 consecutive frames acquired with the flexSPIM, with frame 45 corresponding to stimulus onset. Data from multiple trials were averaged to generate representative activation profiles for the left/right Hb, left/right Rl regions, and left/right Vm regions of the brain. Using ImageJ, regions of interest were manually defined, and the mean fluorescence intensity was measured across all frames, producing time-series traces of average neuronal activity before, during, and after each stimulus.

### Calcium Imaging Data Processing and Spatial Registration

Time-series calcium-imaging data were processed using the open-source CaImAn package (Giovannucci *et al*., 2019) to perform motion correction, signal extraction, and neuronal component identification. Rigid and non-rigid motion correction were applied to minimize motion artifacts and ensure accurate spatial alignment across frames. Following registration, a constrained non-negative matrix factorization (CNMF) model was applied, with key input parameters (e.g., expected neuron diameter, local-correlation threshold, and signal-to-noise cut-offs) empirically optimized to match the imaging resolution and fluorescence dynamics of the dataset. This optimization yielded spatial footprints and corresponding ΔF/F fluorescence traces for individual neuronal components. Model performance was iteratively evaluated through inspection of residual maps and assessment of spatial–temporal consistency.

Neuronal components were selected from anatomically defined regions and depths encompassing the Hb (∼10 µm), Rl region (∼30 µm), and Vm region (∼50 µm). Each region consisted of two bilaterally symmetric areas measuring approximately 50 µm × 50 µm. The Hb regions were located at the most caudal aspect of the brain, the Rl regions were positioned rostrally and laterally, and the Vm regions were situated ventrally and medially, near the midline of the brain. Within each region, ΔF/F fluorescence traces were extracted for all identified neuronal components.

To correct for gradual fluorescence-intensity drift resulting from photobleaching, a rolling mean filter with a window size of 10 frames was applied to stabilize the ΔF/F time series. For each stimulus type, five trials were performed, except for the partial-looming stimulus, which was presented three times. For each trial, 44 frames preceding (∼1min) and 45 frames following (∼1min) the stimulus onset were extracted and averaged across trials, with frame 45 corresponding to the time of stimulus delivery. The time series were then averaged across five trials per fish (three trials for the partial-looming stimulus) for the left and right hemispheric regions of the Hb, Rl, and Vm.

To determine which neuronal components exhibited activation or inhibition in response to the stimulus, the mean ΔF/F values from frames 45-50 (post-stimulus window) were compared with the mean of the preceding 40 frames (baseline). Components showing ΔF/F ≥ 50% relative to baseline were classified as activated, while those with ΔF/F ≤ – 50% were classified as inhibited.

Binary activation maps, showing activated and inhibited components for all fish and stimulus conditions were then registered onto a canonical template fish (three templates: Hb, Rl, and Vm), with activated components represented in red and inhibited components in blue. The canonical template was constructed to represent an “ideal” reference fish and to standardize minor morphological variations across individuals. Registration parameters were manually ascertained using ITK-SNAP, using the midline and the skin-brain boundaries as anatomical landmarks. Using these parameters, each image analyzed was geometrically transformed (cropped, scaled, rotated, and translated) to align with the appropriate canonical template using custom Python scripts incorporating the libraries OpenCV, Matplotlib, and NumPy. The XY coordinates of activated and inhibited components were then extracted from the registered images and plotted onto the canonical template to generate composite activation/inhibition maps showing bilateral regional responses to each stimulus.

### Heating Tg[tiam2a:GFP] Larval Zebrafish using an IR Laser (Figure 9)

Experimental procedures in this section were partially adapted from (Dempsey et al., 2022). To begin, Tg[tiam2a:GFP] zebrafish larvae were embedded in custom behavioral training molds: a 60 mm culture dish filled with 1% agarose dissolved in egg water, with a 10 mm diameter circle cut out from the center. The zebrafish were anesthetized and then embedded in the 10 mm diameter circle with 500 ml agarose + anesthetic (70 ppm MS-222 + 70 ppm Isoflurane (SKU 012ABB-250CC). While the agarose solidified, the zebrafish’s position was parallel to the surface of the chamber, at the top of the agarose surface, and was maintained using a single bristle brush. After 5 min, 10 mL of sterilized husbandry system water was added to the chamber. Using fine forceps, the tail of the zebrafish was carefully freed from the agarose. All free-floating agarose was removed from the dish. The dish was then positioned under the objective of a custom-built behavioral microscope such that the right eye of the zebrafish was directly under where the IR laser spot would be when turned on (NIR RLCO-980-1000-F laser, Roithner Lasertechnik). An Arduino was programmed to deliver the IR heating laser for two seconds, a total of ten times, with an intertrial separation of 2 minutes. After the fish underwent ten trials, it was then fixed in PFA prior to pERK immunohistochemistry.

### pERK Immunohistochemistry

After exposing 7–9 dpf Tg[tiam2a;GFP] larval zebrafish to the IR laser ten times, zebrafish were fixed in 4% PFA diluted in 1x PBS (pH 7.4), 70 ppm MS-222, 70 ppm Isoflurane, covered in aluminum foil, and kept at 4°C overnight. Then, fish were washed 3 times in 1x PBS (pH 7.4) and 0.25% PBST (0.25% Triton X-100 in PBS). After 3x PBST washes, the zebrafish were placed in 150 mM Tris HCl (pH 9) for 5 min at room temperature, followed by 20 minutes at 65°C. Once this was complete, the samples were cooled to room temperature and given 3x PBST washes. Zebrafish were then permeabilized on ice for 45 minutes with 1x PBS, 0.05% Trypsin-EDTA (Sigma, T4049-100ML) diluted in 1x PBS (pH 7.4). After another 3x PBST wash cycle, the zebrafish were placed for 1 hour at room temperature in blocking buffer solution; (1x PBS, 1% bovine serum albumin, 2% DMSO, 2% NGS, 0.25% Triton X-100). Zebrafish were stained with primary rabbit anti-phosphorylated Erk1/2 antibodies at a 1:300 dilution (Cell Signaling, Catalog #4696) in blocking buffer on a rotating platform at 4°C overnight. After 5X PBST washes, the zebrafish were transferred to a blocking buffer solution containing a 1:1000 dilution of Alexa Fluor 568-conjugated goat anti-rabbit secondary antibody (Invitrogen, A-11011) and incubated overnight at 4°C on an orbital shaker in the dark. After 5 PBST washes, samples were moved to PBS solution and imaged on a Zeiss LSM 780 confocal microscope at the Translational Imaging Center at USC.

### Statistical Analysis

Calcium signal curves and heatmaps were calculated using Python 3.12 scripts. Statistical differences for pre– vs post-stimulus AUC values were assessed using one-tailed paired t-tests implemented in the SciPy library, and *p* < 0.05 was considered statistically significant. No statistical methods were used to predetermine sample size.

### Experimental Setup and Stimulation Protocols for Neuronal Activity Imaging in Fish

All fish used in this study were given 90 min to acclimate on the flexSPIM system. Following acclimation, each fish was imaged for 5 min to establish a baseline for neuronal activity before being exposed to stimuli. The total duration of the experiments was ∼15-20 min. After the experiment was finished, the fish was released from the agarose and allowed to rest in a dish for one hour. Only fish that were still alive and healthy after this waiting time were included in the analysis.

### Near-Infrared laser

Once the fish was prepared for imaging, it was positioned such that its right eye was hit by the IR laser (NIR RLCO-980-1000-F laser, Roithner Lasertechnik). While imaging, the IR laser was turned on for 2 seconds before being turned off. This was repeated 2 minutes later for a total of 5 trials.

### Electric shock

Once the fish was prepared for imaging, two 1-pin dual-male jumper wires were secured onto the dive bar such that the pin connectors were positioned on or near both sides of the fish, where the body meets the tail. They were positioned such that when current is applied, it travels between the wires, creating a circuit with the fish in between, causing a small electric shock. While imaging, a 500 mA electric shock was administered to the zebrafish through electrodes connected to a 5 V Arduino pin for 400 ms. This was repeated every 2 minutes, for a total of 5 trials.

### Full-looming Stimulus

Once the fish was prepped for imaging, a red-backlit projector screen was positioned on the right side of the fish. The window on the chamber covered 66° of the fish’s visual field. A small black circle (1° of the fish’s visual field) was maintained throughout the duration of the experiment. At the beginning of each trial, the spot was expanded until it fully covered the entire projection screen (1° to 66° of the visual field) over ms, hence the name “full-looming stimulus”. Following 5 s, the stimulus returned to its original size. This was repeated 2 min later for a total of 5 trials.

### Partial-looming stimulus

Once the fish was prepped for imaging, it was exposed to a stimulus similar to the full-looming stimulus, except that the black circle expanded to 48° over 300 ms, partially covering the visual field. The black circle remained at 48° for 5 seconds before contracting back to its original size of 1° over 300 ms. This process was repeated 2 min later, for a total of 3 trials.

### Tapping

During imaging, the experimenter tapped the side of the chamber with a plastic stick for 5 s. This process was repeated 2 min later, for a total of 5 trials.

### Airhorn

While imaging, an air horn (1.4 oz, Betterboat) sounded a 120-130 dB blast approximately 1 inch above the chamber. The experimenter wore ear protection to prevent hearing damage. This procedure was repeated 2 minutes later, for a total of 5 trials.

### Light

Once the fish was prepared for imaging, a red-backlit projector screen was positioned on the right side of the fish, making the light visible to the fish. A small, removable cardboard card was then placed between the fish and the projector, keeping the fish in complete darkness. During imaging, the card was removed for 5 s to expose the fish to light. After 5 seconds, the card was placed back between the fish and the projector to block the light. This procedure was repeated 2 minutes later, for a total of 5 trials. Activity data was collected for the light-to-dark transition.

## Data Availability

Raw and processed data on Jupyter notebooks containing analysis pipelines are available on https://gin.g-node.org/coltonsm/Zebrafish_Valence_Project. The DOI for this repository will be made public upon being accepted for publication.

## Acknowledgements

We thank M. Jones for microscopy technical support in the Translational Imaging Center, K. Keomanee-Dizon for the design and construction of the SPIM setup, and L. Trinh and E. Carranza Lopez for zebrafish husbandry support. All microscopy was done at the Translational Imaging Center. This work was supported by the National Institute of Mental Health Grant 1UF1NS122082.

