## Supplemental Figures for "Negative-Valence Neurons in the Larval Zebrafish Pallium"

### Supplementary Figures

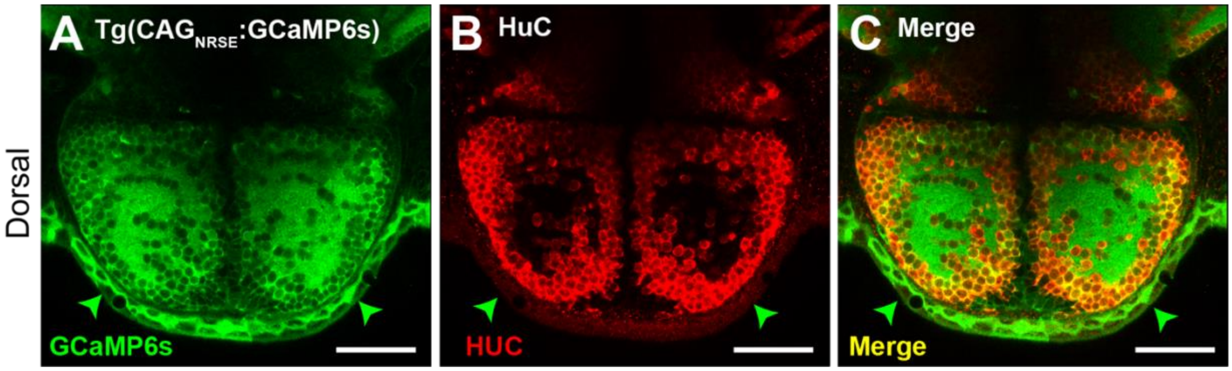


**Figure 1-figure supplement 1. Tg[CAG$_{NRSE}$::PGCaMP6s] expresses GCaMP6s in all neurons**

**(A)** 7 dpf Tg[CAG$_{NRSE}$::GCaMP6s] zebrafish dorsal pallium imaged for GCaMP6s. (**B**) Same zebrafish dorsal pallium immunostained for HuC, which labels all neurons. (**C**) Merge of (A) and (B) shows that essentially all cells labeled for HuC are also labeled for GCaMP6s. Arrowheads point to skin cells expressing GCaMP6s, which were excluded from all analyses.

Scale bar 50 μm.

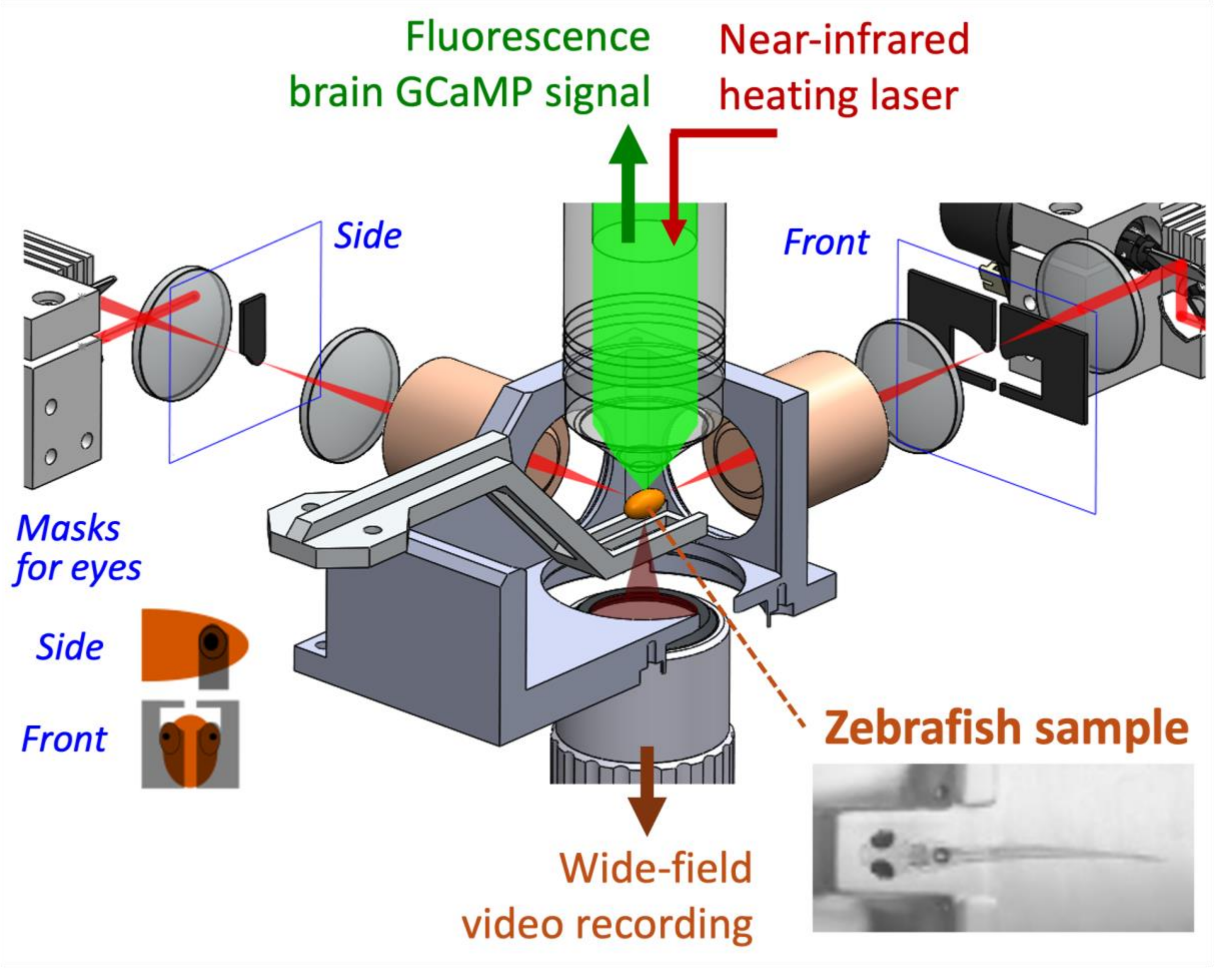


**Figure 1-figure supplement 2. Two-photon light sheet microscopy setup for whole-brain functional imaging**

See (Keomanee-Dizon et al., 2020) for details on design, construction, and operation of the setup. Briefly, the mounted zebrafish sample (depicted as the orange ellipsoid at the center) was positioned inside an aqueous chamber, where excitation light sheets (depicted in red) illuminated the zebrafish's brain from both the front and (left-lateral) side of the animal. Inset at lower right shows a picture of the mounted zebrafish, immobilized in 1.5% low-melt agarose, with its tail freed. Wide-field video recording from the bottom of the sample chamber provided readout of the zebrafish's behavior through its tail movement. The excitation light sheets came from a femtosecond-pulsed near-infrared laser, with wavelength ~920 nm, to generate two-photon-absorption fluorescence from the brain of the zebrafish. The fluorescence signal (depicted in green) was collected by the upright-oriented detection objective and imaged onto a recording camera (not shown). Physical masks made from black-anodized aluminum were positioned along the optical path of the light sheets to prevent direct illumination of the zebrafish eyes. Light from a continuous-wave near-infrared laser, wavelength ~980 nm, was directed toward the zebrafish sample through the detection objective to provide an option for heating stimulation.

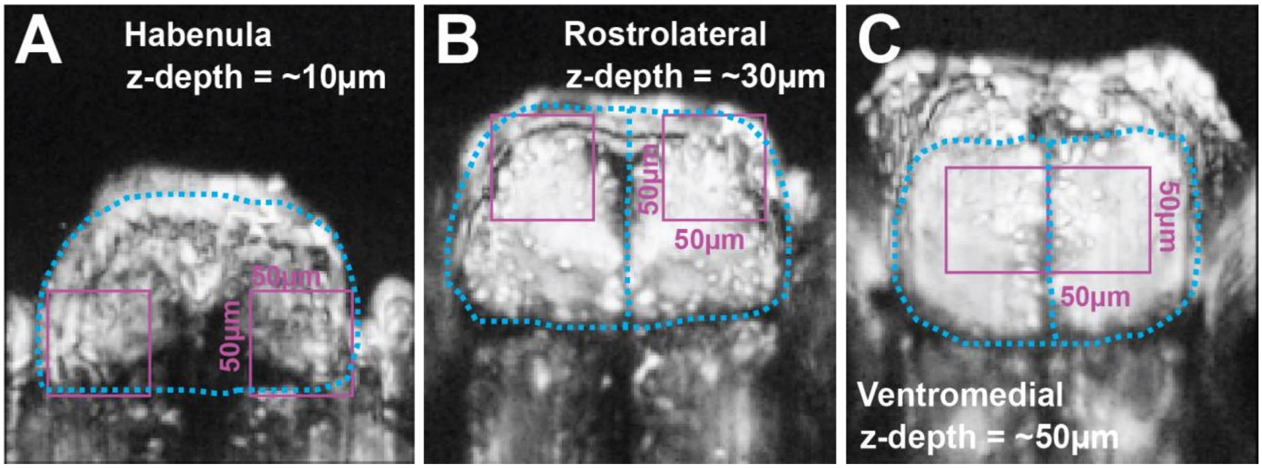


**Figure 1-figure supplement 3. Recording areas**

Recordings were conducted in 50 μm square areas. (**A**) The habenula recordings come from a superficial slice at 10 μm below the surface of the fish brain, 40 μm from the rostral edge, and adjacent to the most lateral edge of the brain. (**B**) The rostrolateral recordings were done at 30 μm below the surface of the brain, adjacent to its rostral edge, and roughly 10 μm from the midline. (**C**) The ventromedial recordings were done at 50 μm below the surface 40 μm from the rostral edge of the brain and adjacent to the midline.

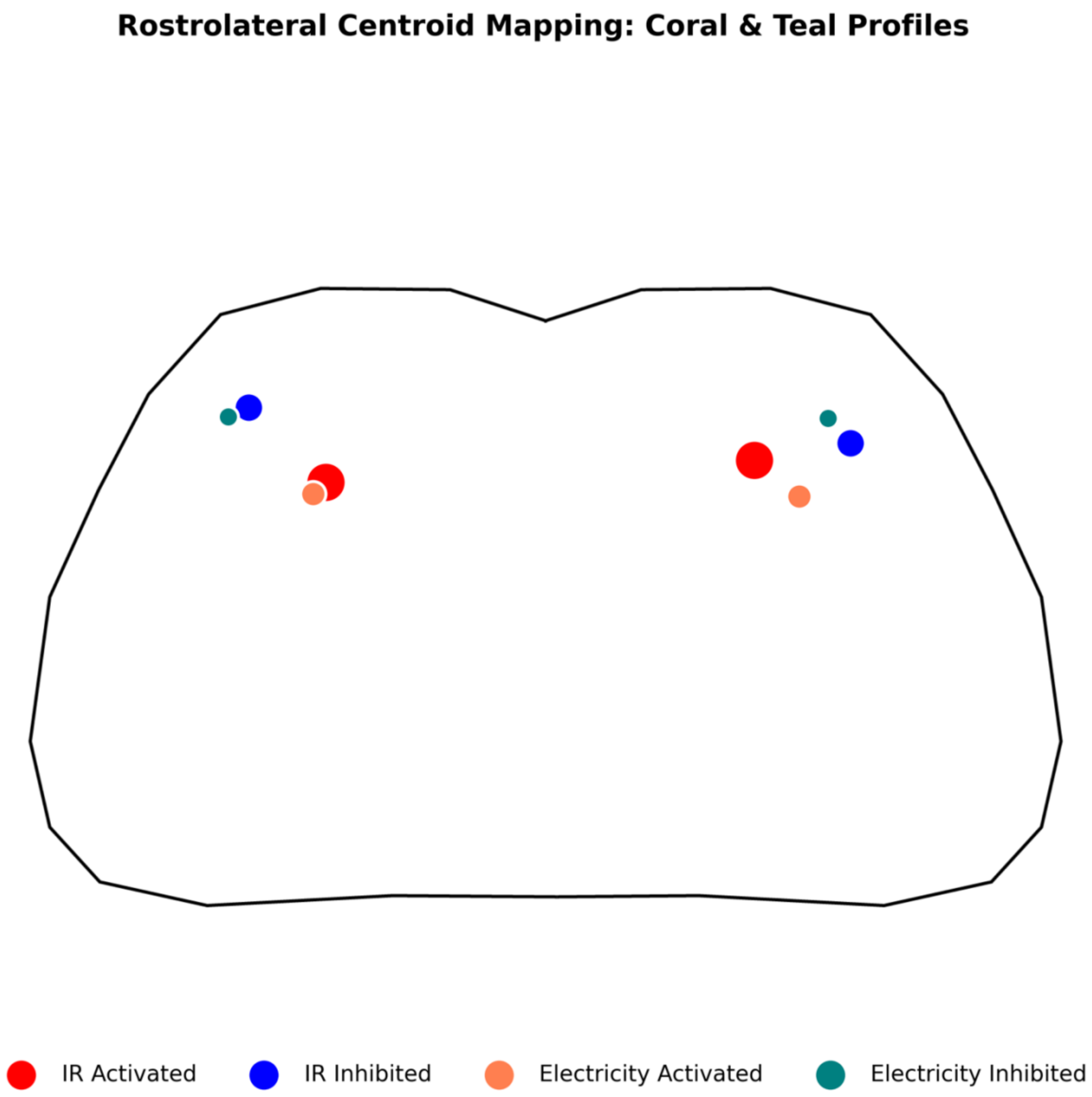


**Figure 2-figure supplement 1.** Overlay of center of mass for excited and inhibited neuronal components in response to IR and electricity. Size of the large dots is proportional to the number of activated or inhibited neuronal components.

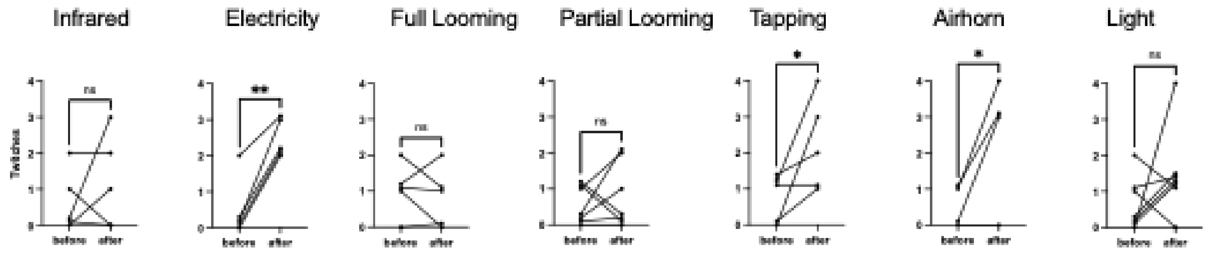


**Figure 8-figure supplement 1** Twitches in Response to Stimuli. Number of twitches of individual fish before and after stimuli. Rates (probability) of twitching before and after were compared with a one-tailed t-test. *, $p < 0.05$; **, $p < 0.01$; ns, $p > 0.05$.
